# Task-related activity in auditory cortex enhances sound representation

**DOI:** 10.1101/2024.08.18.608392

**Authors:** Ana Polterovich, Maciej M. Jankowski, Johannes Niediek, Alex Kazakov, Israel Nelken

**Affiliations:** Edmond and Lily Safra Center for Brain Sciences (ELSC), The Hebrew University of Jerusalem, Jerusalem 91904, Israel; Alexander Silberman Institute of Life Sciences, The Hebrew University of Jerusalem, Jerusalem 91904, Israel; BioTechMed Center, Multimedia Systems Department, Faculty of Electronics, Telecommunications and Informatics, Gdansk University of Technology, Gdansk, Poland; Machine Learning Group, Technische Universität Berlin, Berlin, Germany

**Keywords:** Auditory cortex, rat, behavior, electrophysiology, model, time cells

## Abstract

In auditory-guided tasks, sound presentations often occupy a small fraction of the total task time. We studied here neuronal dynamics that spanned trial duration. Many neurons had large, slow, firing rate modulations, which were not driven by sounds, were larger than the sound evoked responses, and were locked to specific time points during the task, similar to responses of hippocampal time-sensitive neurons. Concurrently, responses to sounds differed between active behavior and passive listening conditions: in the active sessions, the on-going activity just before sound presentations was higher and the responses to target stimuli were weaker but more informative about the task-relevant sounds. We show that the slow firing rate modulations caused the increased on-going activity. Using a model, we demonstrate that higher on-going activity led to more synaptic depression of the cortico-cortical synapses, reducing the tendency to produce population spikes and resulting in weaker but more informative responses.

## Introduction

It has been repeatedly observed that neurons in the auditory cortex encode not only auditory information, but also contextual information^1,2^. Indeed, responses in the auditory cortex are modulated by the sequential context in which sounds are presented^3–9^ and by the task performed by the animals. For example, the engagement of an animal in a task can suppress^10–14^ as well as enhance^11,15^ auditory responses to sounds compared to passive listening. Non-auditory but behaviorally relevant factors also modulate the neural activity of the auditory cortex. Expectations about the timing of an auditory stimulus^16^, involvement in task initiation^17^, attention^18,19^, behavioral choices^20^, behaviorally relevant category^14^ and even expected future reward size^21^ affect the auditory driven responses in auditory cortex. The source of some of these modulations is known: for example, locomotion has diverse effects on sound-evoked responses as well as on on-going activity in primary auditory neurons, and direct projections from M2 may underlie these effects^22,23^. We describe here a new mode of activity in the auditory cortex of freely-moving rats: distinct, slow (over a few seconds) firing rate increases that are locked to trial time, tile trial duration, but are not bound directly to the task-relevant sounds and often precede them. These modulations are different from previously-described modulations of cortical responses, in that they occur at specific moments of the trial, but are not accounted for by sound responses, kinematic variables, or other externally-measured parameters. We conclude that the auditory cortex contains a rich representation of time in trial, from the initiation of a trial and all the way to reward collection. This representation is functionally relevant: it shapes the sound-driven responses, which are weaker, but more informative, during task performance.

## Results

### Behavior

The rats were trained on a combined sound localization-discrimination task in the RIFF (Rat Interactive Foraging Facility), a large arena for studying free behavior and concomitant brain activity in rats^24^. The rats initiated trials by accessing the center of the arena (‘center-crossing’ in the rest of the paper; Fig. 1a). A center-crossing triggered the presentation of a sound, which was then presented every 2 seconds until trial termination. The rat had to go to the interaction area (IA, Methods) indicated by the sound and poke into one of its two ports, receiving a food or water reward (according to the port in which it poked). Poking in a port of another IA aborted the trial, but there were no other consequences and the rat could immediately reinitiate a trial by center-crossing again. Trials were also aborted after 20 s without pokes (‘time out’), but more than 80% of the trials lasted 5 seconds or less (Fig. 1b), with at most 2 sound presentations. There were three types of trials (main task and two controls) that differed in the way sounds indicated the target port (see Methods for details). Since the structure of all trials was identical, we analyze all sessions together.

**Figure 1.**
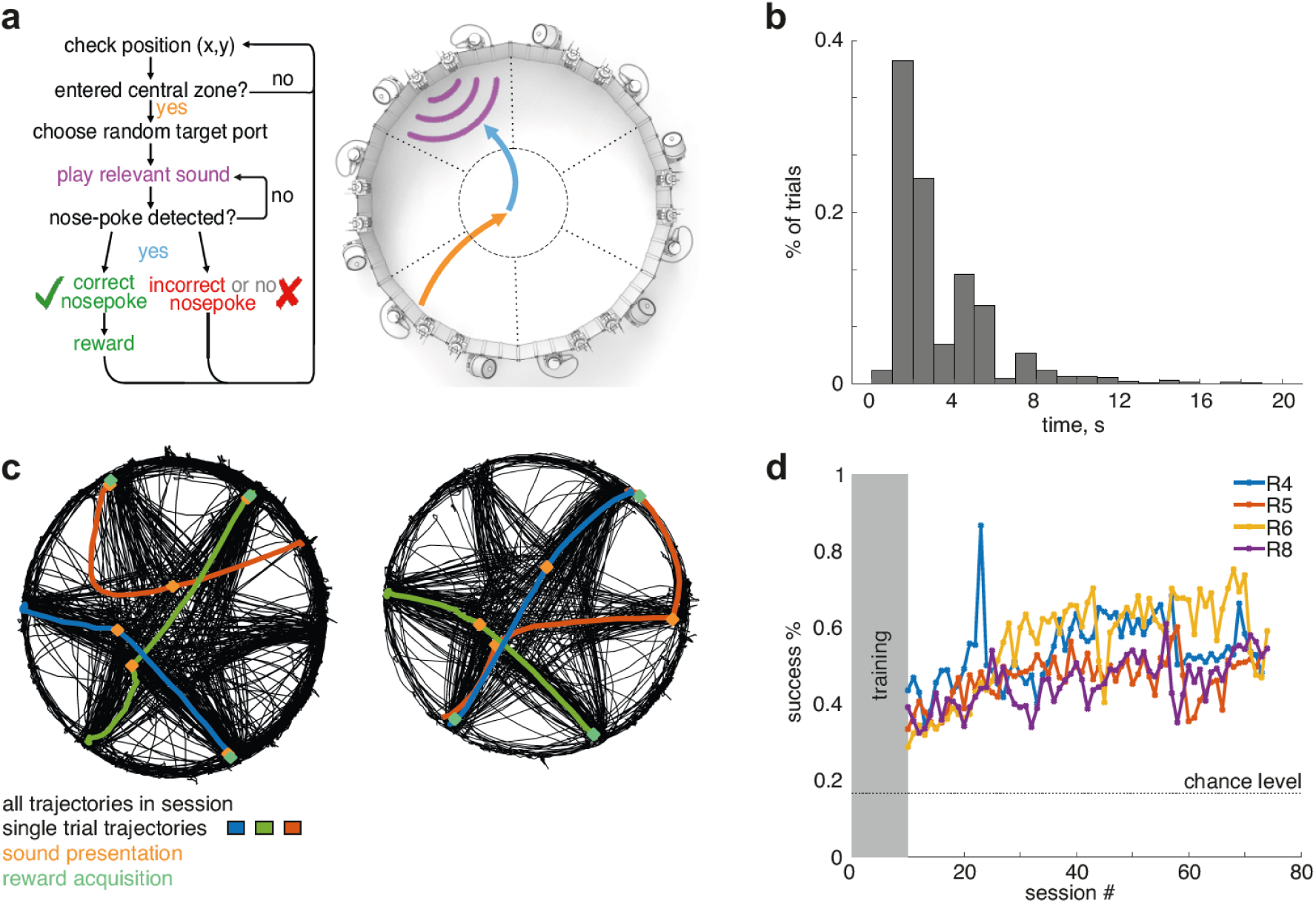
Behavior. a. Scheme of the main task. Flowchart of the experiment real-time loop (left) and the corresponding events in a diagram of the arena (right), plotted with the same color code. b. Distribution of trial durations in all rats. c. Trajectories of 2 example sessions. All trajectories are in black. Highlighted trajectories represent three successful trials (red, green and blue). The location of the rat during sound presentation is depicted in orange, and the location of the rat during reward presentation and acquisition is in light green. d. Success rates in the first 80 behavioral sessions of the full task. Gray area represents the habituation period. Dashed line shows performance at chance level.

Figure 1c illustrates two sessions of the main task (full trajectories in each session in black) with 3 highlighted exemplary successful trials in each session. Already during the first session of the main task (following 10 days of shaping, see Methods), rats performed above chance (Fig. 1d; success rate of 32%±4%, mean±ste, N=4, while random poking would result in a success rate of 17%). The success rate in the last three sessions of the main task was 64%±3% (mean±ste, N=4). Success rates in the control tasks were slightly lower than in the main task, but far above chance level (see Figure S1). These success rates are similar to those reported in the literature for self-initiated complex tasks^25,26^.

We study here two types of neural activity: the sensory responses to the task-relevant sounds during active behavior and during passive listening, and a new mode of activity in auditory cortex that we identified during behavior, consisting of large, slow firing rate modulations that were task related. In the following sections (1) we describe the large, slow firing rate modulations that were locked to task events but often not driven by sounds or other externally-measured variables; (2) we show that sensory responses to the task-relevant sounds were weaker, but more informative, during active task performance than during passive listening; and (3) using a model, we show that the slow firing rate modulations can account for the differences between neuronal responses in the active and passive conditions.

### Large, slow modulations in firing rates locked to trial events

The main novel finding reported here is an activity mode in the auditory cortex that consists of large, slow firing rate modulations that were locked to trial events, lasted a second or more, and were not evoked by the task-relevant sounds. Figure 2 depicts several examples of such events in 4 units from 3 rats. In each panel, the average firing rate of one unit over all trials is depicted in black. Concomitant average running velocity (green) and sound level (derived from the signal recorded by the microphone on the head of the rat, magenta) are displayed below. Firing rates at all individual trials are displayed at the bottom of each panel.

**Figure 2.**
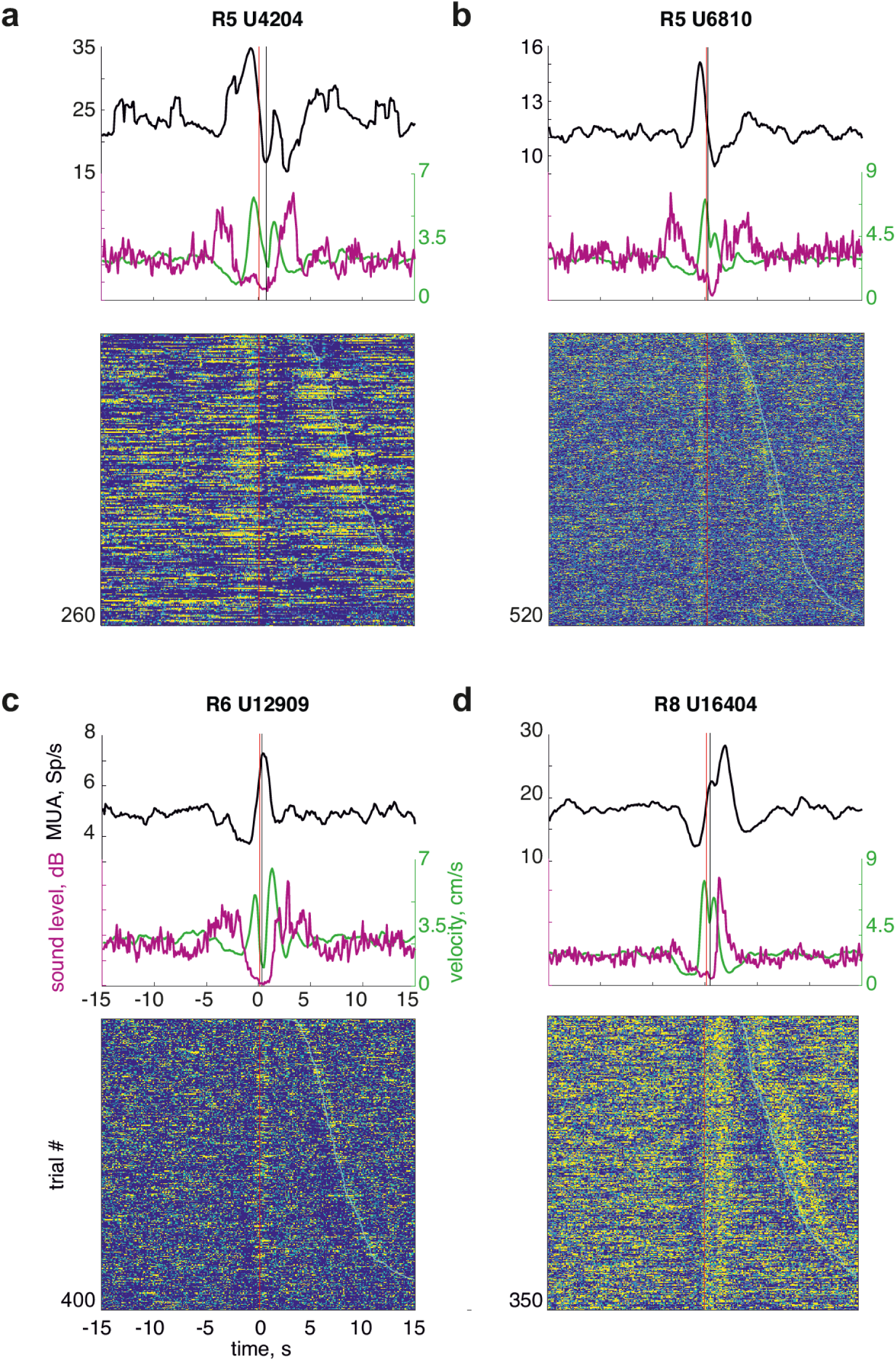
Large, slow firing rate modulations locked to trial events. a. Top: Mean firing rate of a unit (black). The concurrent mean sound level (in arbitrary units, magenta) and mean velocity (green) are presented below. The time axis spans the range from 15 seconds before center-crossing to 15 seconds after center-crossing. The red vertical line indicates the time of center crossing; the black vertical line indicates the time of sound onset. Bottom of the panel: Trial by trial firing rate of the same unit, sorted by the latency to the center-crossing of the following trial. Yellow indicates higher firing rate, blue indicates lower firing rate. The time of center-crossing is indicated by the red line, and the center- crossing of the following trial is indicated by the cyan line. The number of trials is indicated in the bottom left corner. b. c. d. Same as (a), for three other units.

In Fig. 2a, the unit increased its firing rate a few seconds before center crossing, preceding the increase in velocity (green trace) associated with the movement towards the center of the arena. This event lasted 3-4 s, and can be clearly seen in single trials (Fig. 2a, bottom). In Fig. 2b, a similar firing rate event occurred, peaking just before center crossing. The firing rate modulation didn’t match the temporal profile of changes in average sound level, nor that of the average velocity of the rat during the session in which this unit was recorded.

Different units could show such slow transient modulations of their firing rate at different times during the trial. In Fig. 2c, the firing rate peaked shortly after center crossing, but in Fig. 2d it peaked almost a second after the presentation of the task-relevant sound. Importantly, in both Figs. 2c and 2d, the firing rate started increasing before center crossing (and therefore before sound presentation).

Such slow firing rate modulations were a common occurrence. For many neurons, the major firing rate peak occurred within a few seconds before or after sound presentation around center crossing. In Fig. 3a, the normalized firing rates of all the neurons recorded from each rat are arranged by the time of the maximum firing rate within an interval of ±5 s around the center crossing. These peaks spanned the time course of the trial, from a few seconds before center crossing (when the rats started running towards the center of the arena) to a few seconds after center crossing, when they collected the reward. They were largely independent of the position of the rat when it initiated the trial and of the sound that was presented to the rat: Fig. S2 depicts the firing rates of 4 example units, with trials averaged by area at trial initiation or by sound identity, as well as the average over all trials. We compared the temporal variation of the overall mean over all trials (modulation energy) with the deviations (mean squared differences) of the area-specific trial averages from their overall mean. The deviations were smaller than the modulation energy for 92% (dividing by area) and 95% (dividing by sound) of the units.

**Figure 3.**
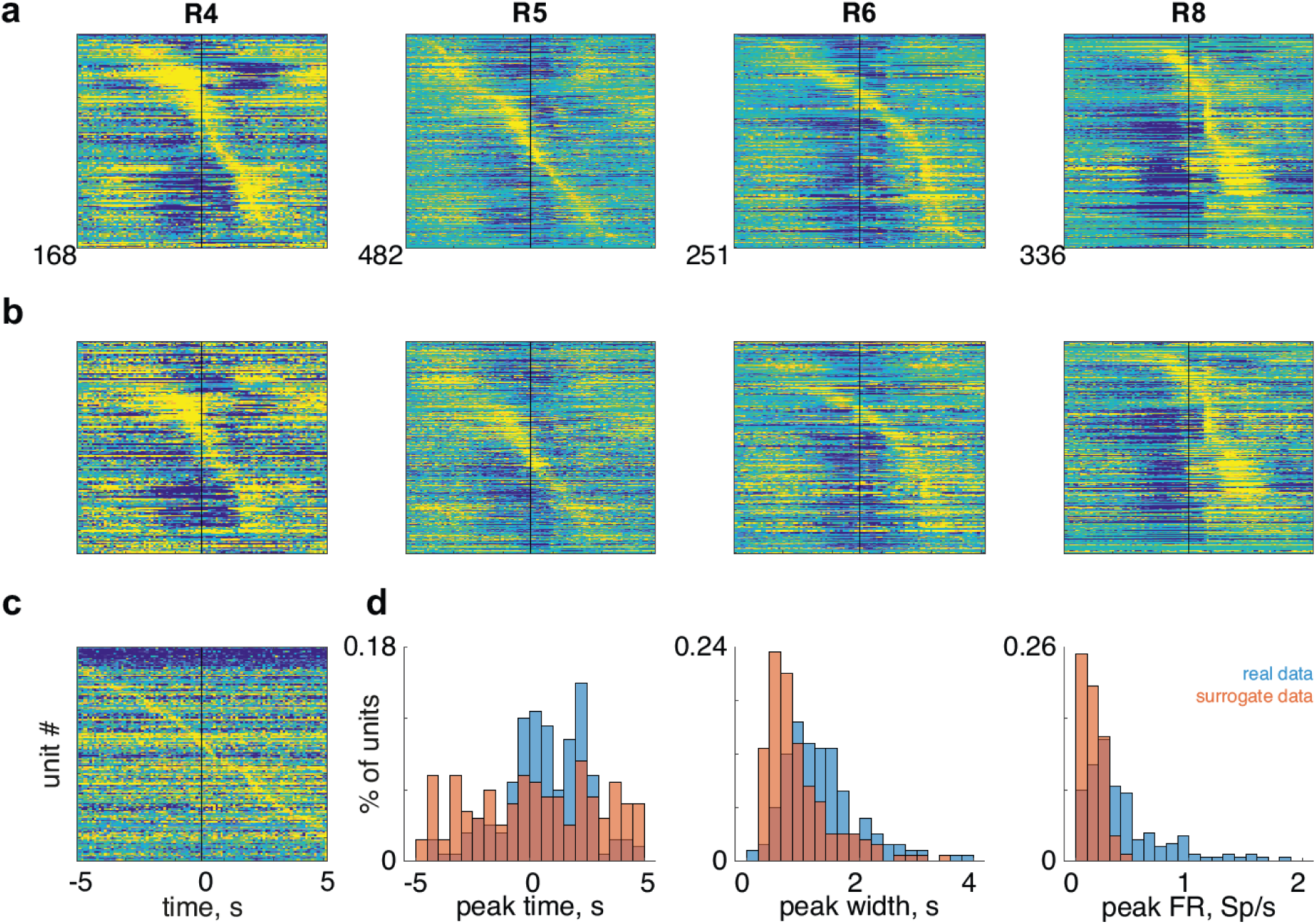
Population representation of time in trial. a. Normalized mean firing rates of all units in four rats sorted by the time of the maximum firing rate (from 5 seconds before to 5 seconds after center-crossing). Yellow indicates high firing rate, blue indicates low firing rate. The black vertical line indicates the time of center-crossing. b. Firing rates averaged over the even trials of each unit, sorted by the time of the maxima of the average over the odd trials. Color code same as in a. c. Mean firing rates around uniformly distributed surrogate center crossing times for rat 4, ordered by maximum times. Same color code as in (a). d. Distribution of peak time (left), width (middle) and height (right) of the maximal firing rates of the real (blue) and surrogate (red) data for rat 4.

To confirm the trial-by-trial consistency of the firing rate modulations, we divided the trials of each session into odd and even trial sets. Figure 3b depicts the firing rates of the units, averaged over the even trials, ordered according to the times of the peaks of the average firing rates over the odd trials (Fig S3a shows the averages of the even trials and of the odd trials separately for every rat). The times of the maxima of the modulations in the odd and even trials for each unit were highly correlated (Fig. S3b; for each rat, χ^2^(16) > 98, p < 10^-13^, see Table S1 for individual animals). As a second control, we selected uniformly distributed surrogate center crossing times within the same sessions and used them to compute firing rate traces for each unit. Figure 3c shows these surrogate traces for one rat (R4), ordered by the time of their peak (data for all rats in Fig S3c). The peaks found in the real data were more concentrated around the center crossing than in the surrogate data (Fig 4d, left panel, two sample Kolmogorov-Smirnov tests, p < 10^-3^ in all cases); the peaks were also wider and higher for the real data than for the surrogate data (Fig 3d; width: middle panel; two-sample t-test for each rat separately, p < 10^-7^ in all cases; height: right panel, two-sample t-test for each rat separately, p < 10^-3^ in all cases, see Table S2 and Fig. S3d for full details). Finally, there was no correlation between the times of the maxima of the modulations when averaging the surrogate data separately over the odd and the even trials (χ^2^(16) < 21, p > 0.2, Table S3 and Fig. S3e-f).

**Figure 4.**
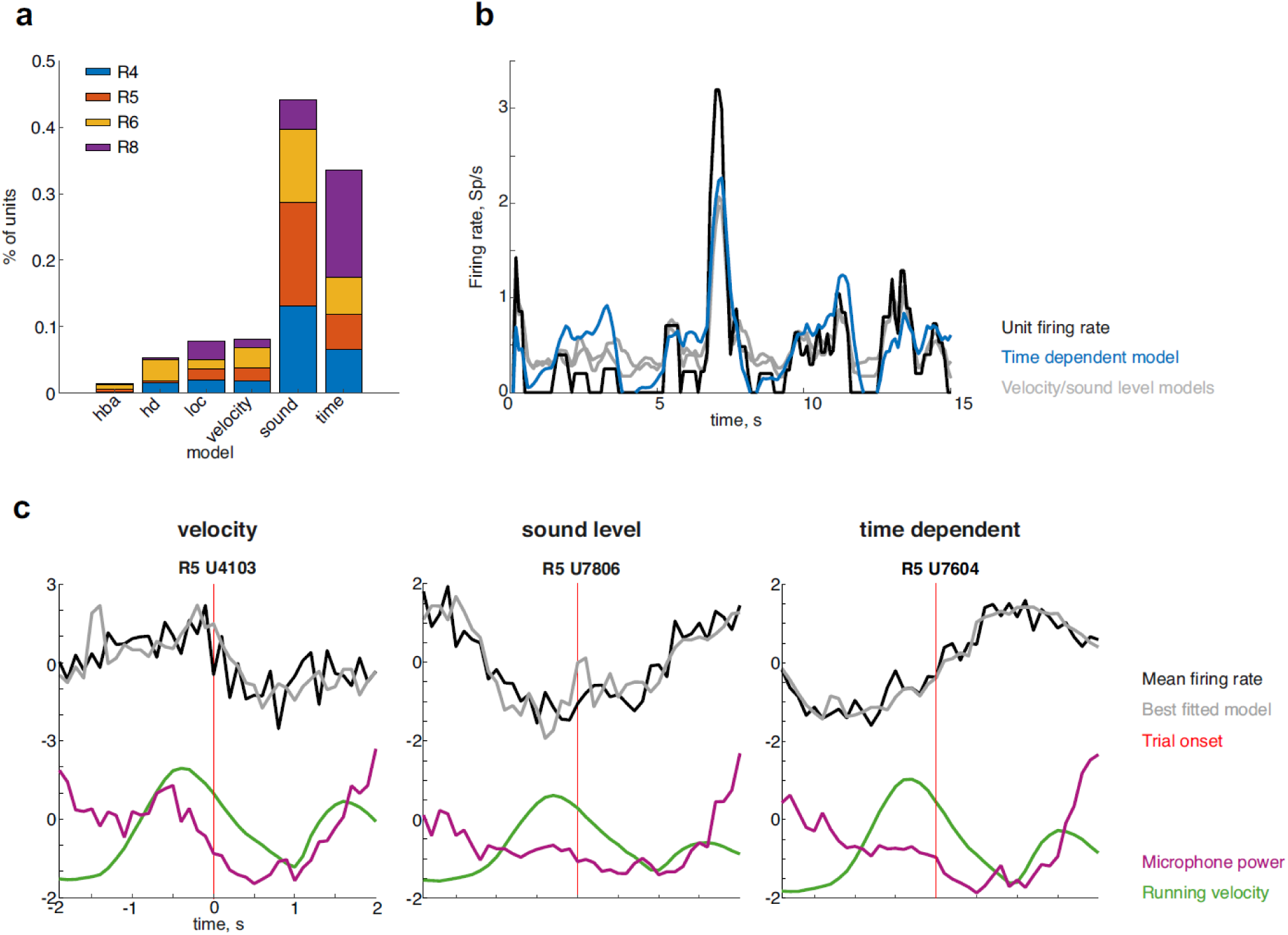
Different variables explain the behavior of the units. a. The distribution of the best explanatory variable for each unit. Rats are represented by colors. HBA: head-body angle; HD: head direction; LOC: location. b. Firing rate during 15 seconds of an example unit (black) best explained by the time dependent model. In blue is the fit of the time model, and in gray the fit of the sound level and running velocity models. c. Z-scored mean firing rates around center-crossing of three example units in black, each fitted with its best model (in gray, from left to right: velocity, sound level and time). Below are the concurrent mean sound level (magenta) and mean running velocity (green). The red vertical line is the time of center-crossing.

Given the above, we determined for each neuron the time of the peak firing rate during the interval between −5 s and 5 s around the center crossing for the odd trials, and we refer to the firing rate transient around this maximum, calculated over the even trials, as the ‘slow firing rate modulation’ for that neuron.

In the examples in Fig. 2, the temporal profile of the firing rate followed neither changes in velocity nor changes in sound level (recorded by the microphone on the head-stage): it is possible to find moments of high firing rate and high velocity as well as moments of high firing rate and low velocity, and similarly for the sound level. To study rigorously the potential causes for the slow firing rate modulations, we tested the ability of five variables to account for the moment-by-moment changes in firing rates (see Methods): sound level, rat velocity, rat location, head direction, and the angle between the head and the body of the animal. We compared the resulting models with a sixth one, in which the firing rate depended on the time in trial (relative to the center crossing). We modeled the firing rates over a total of 4 seconds, starting from 2 s before center crossing and ending at 2 s after center crossing, ensuring that data from successive trials would only minimally overlap. The models were fitted for all units of each rat separately using linear mixed-effects models, in which each parameter had a ‘fixed’ contribution (accounting for the mean firing rate over all units for that rat) and a ‘random’ contribution that was unit-specific. Since the models contained non-overlapping sets of explanatory variables with some variation in their dimensionality, we compared models using the Akaike Information Criterion (AIC; see Methods for details). For all rats, the two best explanatory variables were sound level and time in trial. For three of the rats, the sound level model was the best and for one rat the time in trial model was the best (Table S4).

The models also provided fits for each unit individually. For every unit, we looked for the model that provided the best fit (largest likelihood, see Statistical analysis section for details, and Fig 4a for a summary of the results). The model that provided the largest number of best fits was the sound level model (45%), followed closely by the time in trial model (at 34%). Substantially fewer units had best fit by the velocity model (8%), the location model (8%), head direction model (4%) and head-body angle model (1%). Thus, while a plurality of the neurons were, as expected, sound-sensitive, for about a third of the units, the best explanatory variable for the slow modulations in their firing rates was time in trial.

Figure 4b depicts 15 seconds of the activity (black) of an example unit whose responses were best explained by their dependence on the time in trial. The fitted firing rate from the model is depicted in blue, and the fits derived from the velocity and sound level models are plotted in gray, illustrating the better explanatory power of time in trial relative to the other variables. Figure 4c depicts the average neural activity of three units around the time of center crossing. In Fig. 4c (left), velocity provided the best fit for the firing rate modulations, and indeed the average firing rate is clearly correlated with the changes in average velocity during the same time. In Fig. 4c (center), the sound level provided the best fit, and the modulation of the average firing rate was correlated with that of the average sound level. In Fig. 4c (right) the unit is best fitted by the time-dependent model (same unit as in Fig. 4b). In this example, the firing rate modulations did not correlate well with either velocity or sound level.

### Responses to task-relevant sounds differed between passive and active listening

Figure 5a depicts the oscillogram and spectrogram of one of the task-relevant sounds (“da”, marked by the color red in the other panels of Fig. 5). The other five task-relevant sounds are displayed in Fig. S4. Responsiveness was similar in passive sessions (74% of all units had significant responses to at least one sound, 48%-94%, N=4 rats) and in the active sessions (69% of the units had significant responses to at least one sound, 47%-78%, N=4). However, there were striking differences in the activity of the same neurons in the active and passive conditions. Figure 5b shows a comparison between the spike counts in the active and passive conditions for all units of one rat (see Fig. S5 for the data of all rats). Ongoing firing rates during the 200 ms preceding sound onset were significantly higher in the active sessions compared to the passive sessions (linear mixed effects model, Table S5; significant effect of condition, F(1,12262)=238, p=3.5*10^-53^; Fig. 5c, left). At the same time, auditory responses to the task-relevant stimuli were significantly weaker in the active sessions compared to the passive sessions (Table S6; significant effect of condition, F(1,12262) = 241, p = 6.2*10^-54^; Fig. 5c, right).

**Figure 5.**
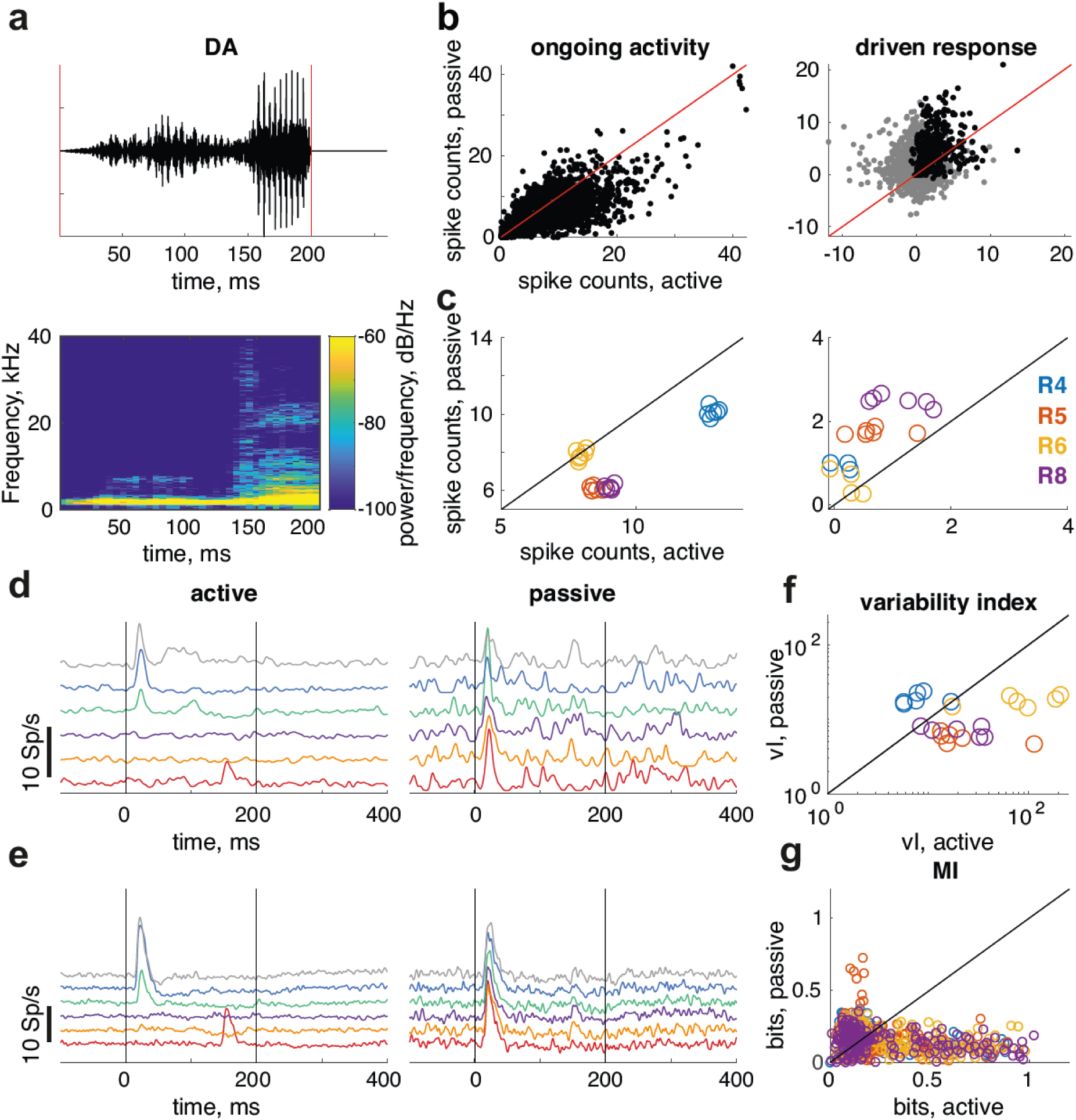
Responses to the task-relevant sounds. a. Oscillogram (top) and spectrogram (bottom) of the word “da”, one of the 6 stimuli presented to the rats. b. Mean spike counts of the units from rat 5 during active compared to passive sessions. Each dot is one unit. Left: On-going activity in the 200 ms preceding sound onset. Right: responses during sound presentation (0-200 ms after sound onset). Black dots indicate sound responses that were significant in both passive and active conditions. c. Mean responses to each stimulus for each rat in passive compared to active sessions. Each circle is the mean response to one of the stimuli in all units of the same rat. Each color indicates a different rat. Left: on-going activity (200-0 ms before sound onset), Right: sound driven response (0-200 ms after sound onset).Yellow circles summarize the data presented in (b). d. Mean responses of one unit in rat 6 to each one of the 6 stimuli. Each color indicates one stimulus. Left: active sessions. Right: passive sessions. e. Mean responses of all units in rat 6 to each one of the task-relevant stimuli. Each color indicates one stimulus. Left: active sessions. Right: passive sessions. f. Variability index for each stimulus in active compared to passive sessions, on a logarithmic scale. Each circle indicates one sound. Each color corresponds to a different rat (same color code as in c). g. Mutual information between response and stimulus identity in active compared to passive sessions. Each circle indicates one unit. Colors correspond to different rats.

The temporal patterns of the responses to the same sound also differed between the active and passive sessions. They tended to be richer and more distinct in the active condition. The responses of one unit to all six task-relevant sounds are illustrated in Fig. 5d. For example, while in the passive condition the response to “da” (red in Fig. 5d) occurred shortly after sound onset, the response in the active condition was elicited later, at the voice onset time^27^. Figure 5e depicts the mean response of all units in rat 6 in the active and the passive conditions (left and right respectively). In the passive condition the responses to all sounds were evoked by sound onset. In contrast, in the active condition the responses to the different stimuli differed in their strength as well as in their temporal patterns. The variability index (Fig. 5f) compares the temporal response pattern, averaged over all sounds, with the responses to each individual sound, so that it is small when the responses are similar to each other, and large when they differ (see Methods). The variability index was indeed larger in the active compared to the passive condition for most rats and sounds. The higher variability was beneficial for coding sound identity: the mutual information (MI) between sound identity and the single-trial responses tended to be larger in the active condition compared to the passive condition (Fig. 5g; Table S7; main effect of condition, F(1,1700) = 107, p = 2.2*10^-24^).

### The large, slow firing rate modulations shape the responses to sounds in the active state

We documented two differences between the neural activity in the active and passive conditions: the on-going activity was higher in the active condition, while the neural responses were lower but more informative. We argue here that both effects are likely a consequence of the slow firing rate modulations that occur in the active condition.

To start with, the number of spikes involved with the slow firing rate modulations were substantially larger than the evoked responses as shown in Fig. 6a (Table S8; significant effect of type, F(1,2472)=41.8, p=1.2*10^-10^). Thus, the firing rate modulations could exert a powerful effect on the sensory responses.

**Figure 6.**
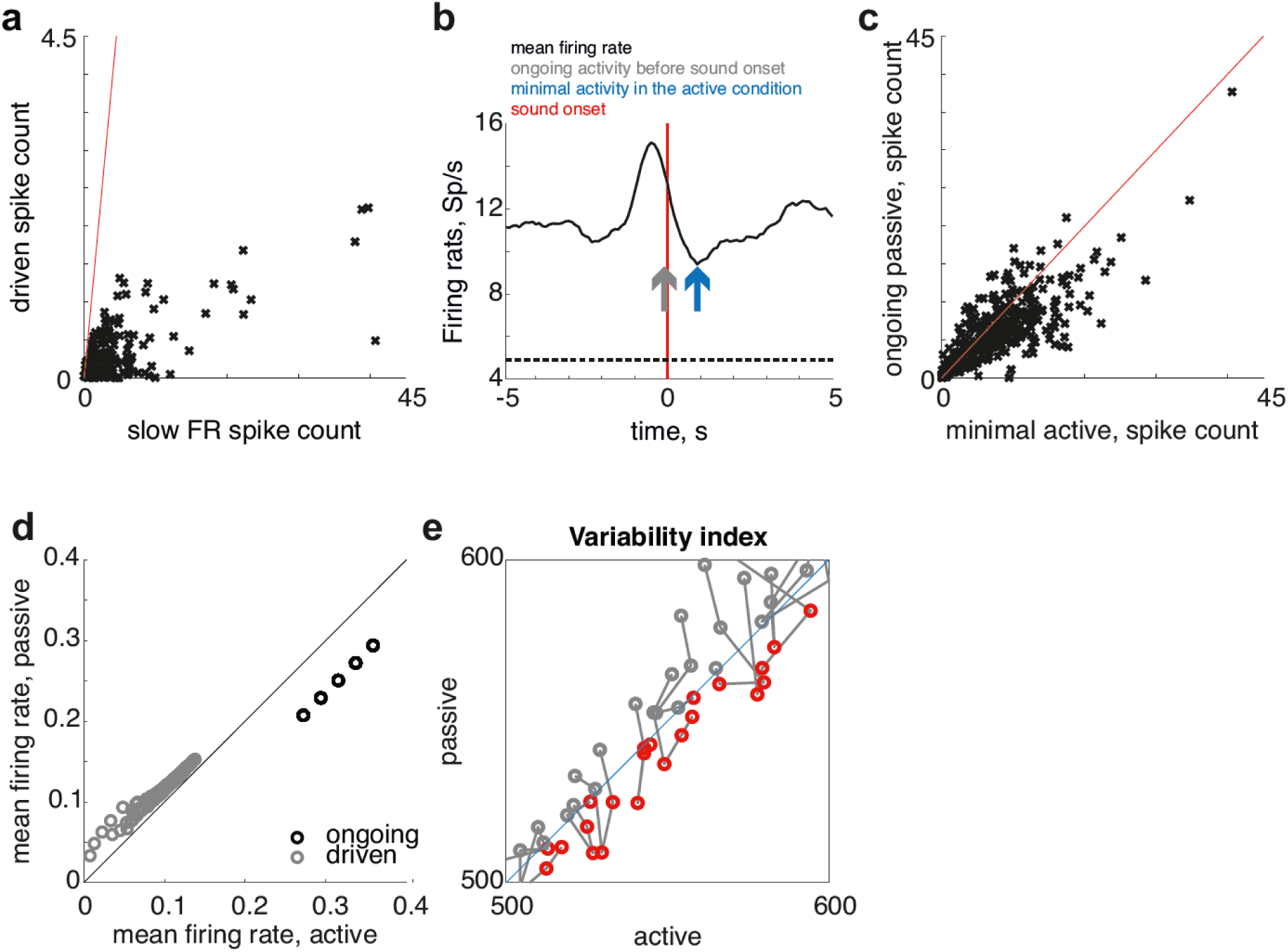
Large, slow firing rate modulations account for the differences between the responses to the task-relevant sounds in passive and active sessions. a. Mean spike count during the slow firing rate modulation compared to sound driven response in R5. Each dot depicts one unit. b. Definitions of “on-going activity” and “minimal activity”: on-going activity (gray arrow) was the average firing rate in the interval 200-0 ms before sound onset (red vertical line). The minimal activity (blue arrow) was the minimal firing rate in the ±5 second range around sound onset. The dashed line indicates on-going activity in the passive condition for this unit. c. On-going activity in the passive sessions compared to the minimal activity in the active session for all units from R5. Each dot depicts one unit. d. Model driven (gray) and on-going (black) firing rates in the active compared to the passive conditions. e. Variability index of the model output in the active compared to the passive conditions. Red circles indicate cases where the variability was larger for the active condition. Circles that are joined by a line correspond to simulations with the same sound level that differed in the on-going passive (and therefore also active) firing rates.

Next, we show that the slow firing rate modulations account for much of the increase in on-going firing rates in the active state relative to the passive state. Figure 6b shows the average activity of an example unit during behavior. For comparison, the on-going firing rate just preceding sound onset during the corresponding passive session is marked by a dashed line. The passive on-going rate may be usefully compared with two values along this trace. The first is the on-going firing rate just preceding sound onset, marked by a gray arrow. As shown before (Fig. 5c), the passive on-going rate is lower on average than the active on-going rate just preceding sound onset. Figure 6b provides a simple explanation for that observation: sound presentations in the active sessions may occur while the slow firing modulation unfolds, raising the firing rate of the neuron. The second possible comparison value is the minimum firing rate of the unit in the displayed time window (marked by a blue arrow). Indeed, while the minimal activity in the active condition was somewhat larger than the on-going passive activity, the difference failed to reach significance (Table S9, post hoc comparisons between the two conditions, minimal active > ongoing passive, F(1,3114)=3.81, p=0.051). More importantly, the minimum firing rate in the active condition was highly correlated with the on-going passive firing rate: the correlation coefficients ranged between 0.47 and 0.77 for the four rats (Fig. 6c, p < 10^-10^, see Table S10 for individual rats). We conclude that the difference between the firing rates preceding sound presentations in the active and in the passive conditions (as shown in Fig. 5c) is largely due to the slow firing rate modulations that increase the on-going firing rate in the active condition above their minimum value.

The increase in the on-going activity prior to sound presentations, which is a consequence of the slow firing rate modulations, may have significant repercussions on the sensory responses in the active condition. Indeed, cortico-cortical synapses in the auditory cortex show a substantial amount of synaptic depression^28,29^, so that increases in cortical firing rates in the active condition are expected to translate into more depressed cortico-cortical synapses in the active condition, and therefore to smaller sensory responses in the active condition.

To quantify this intuition, we used the model of Yarden and Nelken^30^ as our starting point (See Methods for details). This model, in which synaptic activation is followed by depression that recovers exponentially^30,31^ produces ‘population spikes’. These are large, coordinated events where multiple neurons fire in close proximity. Population spikes may mask differences between stimuli. They depend on the background level of neural activity and tend to disappear at high levels of background neural activity because of the synaptic depression of the cortico-cortical synapses. When they don’t occur, the network may show increased sensitivity to fine features of its inputs^31^. We used the model to show that a similar mechanism may operate here too.

The model is built from densely connected networks that represent cortical columns, with sparser inter-columnar connections. The neurons within each column receive their input from similar frequency channels, with some variability (to mimic the imprecise tonotopy typical of the rodent auditory cortex^32^). For the purpose of the simulation, the input to each column was computed by feeding the six sounds used in the experiments through an auditory model of the peripheral auditory system^33^. The auditory representation was then low-pass filtered to represent the slowing down of the responses along the ascending auditory system^34^, and high-pass filtered to make the responses more tonic, as typical of higher stations of the auditory system^35^.

We initially used the exact same network parameters as in Yarden and Nelken^30^. With these parameters, the responses of the network consisted of onset responses for most sounds at most frequency channels. These responses resemble the responses of the cortical neurons in the passive condition (as in Fig. 5b and Fig. S5b).

Because of the strong dependence of synaptic depression on the firing rates of the neurons, the network can be controlled by modifying the baseline firing rates of its neurons. To simulate activity in a passive and in an active state, we compared responses simulated at one baseline firing rate to the responses simulated with a baseline firing rate that was 30% higher, similar to the average increase in on-going rate in the active condition just preceding sound onsets (Fig. 5f and Fig. S5c). The on-going firing rates increased in all variants of the simulation (Fig. 6d, black points) while evoked neuronal responses decreased (Fig. 6d, gray points; each point represents the response to one of the six sounds, in one pair of firing rate conditions), as we observed in the active condition (Fig. 5g). Thus, the model easily reproduced the observation that higher on-going rates are associated with weaker sensory responses.

Finally, Fig. 6e compares the variability index (as in Fig. 5d) of the model responses in the active and passive conditions. The variability index was computed after averaging the responses to each of the six sounds over all neurons in each baseline firing rate condition. Points that are joined by a line correspond to simulations with the same sound level that differed in the baseline firing rates (one value for simulating the passive state, a 30% higher value for simulating the active state). Red circles depict cases in which the variability index in the active condition was higher than in the passive condition, as we observed in the data. Figure 6e shows that there is a wide range of baseline firing rates and sound amplitudes for which the variability index in the active state is higher than in the passive state. Thus, without fine tuning, in a validated model of auditory cortex, simple changes in the on-going activity preceding sound onset are sufficient to reproduce the observed changes in firing rates and richness of response patterns.

## Discussion

The main result of this paper is the demonstration, in the auditory cortex of behaving rats, of large, slow firing rate modulations that were locked to task events but often were not evoked by sounds. We showed that these modulations occurred in many neurons, and covered the relevant time period between the initiation of the trial by the rat (a couple of seconds before center crossing) to reward (a couple of seconds after center crossing). The number of spikes that composed these slow firing rate modulations was substantially larger than those emitted as part of the responses to the task-relevant sounds. We argue that this new mode of activity had an important function: it shaped the sensory responses and made them more informative when the rats were actively performing the task, relative to periods when the rats listened passively to the same sounds.

These results are consistent with a number of previous studies. A reduction in the responses during the performance of a task has been shown in ferrets^12–14^ and in mice^10^. In these studies, sensory responses were weaker during active behavior compared to passive listening. Otazu et al.^10^ traced this difference to higher ongoing rates in the auditory thalamus, and using a model similar to ours argued that such an increase would cause a depression of the thalamo-cortical synapses leading to weaker responses. Bagur et al.^13^ as well as Yao et al.^36^ found that responses in the auditory cortex were more informative during engagement in a task than during passive listening.

We argue that the reduction in the task-related responses shown here is a consequence of the higher ongoing activity just before stimulus onset in the active compared to the passive conditions. These results are in concordance with the findings of Bagur et al.^13^, who showed that the ongoing activity during the stimulus presentation was enough to explain the differences in responses to target and reference stimuli between the passive and active conditions. These results are somewhat different from those of Otazu et al.^10^ who found that the ongoing firing rates in cortex were similar in the engaged and passive conditions. The difference between Otazu et al.^10^ and the current study could be due to species difference (mice vs. rats), the behavioral setup, or the task.

An alternative mechanism that accounts for the smaller but more variable responses during the active behavior has been suggested^11^. They showed that in mice, inhibitory interneurons in auditory cortex were upregulated in an active task compared to passive listening, causing some suppression of the responses of excitatory neurons to auditory stimuli when they were behaviorally meaningful (SOM+ and PV+ neurons) and upregulating the on-going firing rates during the active task (VIP+ neurons). This mechanism may contribute to our results as well, although its interactions with the slow modulations are unknown.

The slow firing rate modulations we describe here are remarkably similar to the activity of ‘time neurons’ that have been found in the hippocampus^37–41^, medial entorhinal cortex^42^, striatum^43,44^ and even prefrontal cortex^44,45^. A time neuron fires at a specific time point during well-practiced tasks; these times are characteristic to the neuron, but vary between neurons and may span the whole duration of the task.

In our task, rats were required to repeat a well-trained behavioral pattern (access the center of the arena, make a decision, and then run to an interaction area in order to collect reward). The times of the firing rate modulations were consistent relative to the start of trial on a trial-by-trial basis. They were largely independent of the location of the rat at trial initiation and therefore also of the running path to and from the center of the arena. While some of the modulations of a plurality of the neurons could be explained by sounds that occurred in the arena (most of them likely self-generated), for about a third of the units the best model for their firing rate modulations depended on time in trial. These are the properties that are strongly reminiscent of ‘time neurons’ in the hippocampal formation^37–42^. Since we recorded only from well-trained rats, we do not know whether these neurons existed before learning, or developed their special properties after the rats learned the task.

The time-sensitive activity could be locally generated through the dynamics of neural interactions and population activity in the auditory cortex itself^11^; we recorded only about 10 neurons in each session, limiting our ability to study the dynamics. Alternatively, time-sensitive inputs could shape this activity. One potential source for such activity is the prefrontal cortex (PFC). The mPFC has both inputs from and outputs to the auditory cortex^46^, and has been shown to have ‘time neurons’^44,45^. Another possible source of time-sensitivity to neurons in the auditory cortex is the hippocampus itself. Hippocampal activity can be relayed to the auditory cortex through the medial entorhinal cortex^47^.

This interpretation of the data is somewhat limited by the experimental constraints. We cannot reliably identify cell types and layers in our data, and so do not know the exact cellular identity of neurons that had such responses. There could also be other external factors that we failed to consider, and that could provide an explanation for the slow firing rate modulations. However, the two major known drivers of activity in the auditory cortex are sounds and locomotion^22,23^, and we tested both carefully.

The finding of time-related responses in the auditory cortex, consisting of large firing rate modulations that tile trial duration, highlights the multifaceted role of the auditory cortex in complex auditory guided behavior. In addition to coding sounds within their context^1–9^, we show here that the auditory cortex literally keeps time.

## STAR Methods

### Resource availability

#### Lead Contact

Further information and requests for resources and reagents should be directed to and will be fulfilled by the lead contact, Israel Nelken.

#### Materials availability

This study did not generate new unique reagents.

#### Data and code availability

● The MATLAB code to create the figures and a sample of the processed data have been deposited at Figshare and are publicly available as of the date of publication at https://doi.org/10.6084/m9.figshare.26310223
● Any additional information required to reanalyze the data reported in this paper is available from the lead contact upon request.

#### Experimental model and subject details

The experiments were carried out in accordance with the regulations of the ethics committee of The Hebrew University of Jerusalem. The Hebrew University of Jerusalem is an Association for Assessment and Accreditation of Laboratory Animal Care (AAALAC) accredited institution. Four adult female Sabra rats (Envigo LTD, Israel; weight: at least 200 gr) were used for the experiments described here. All efforts were taken to create a low-stress, rat-friendly living environment enabling experimental animals to freely express their innate behaviors. At their arrival, the rats were housed in the same SPF room in which the experimental setup was situated and where all experiments took place. The animals were housed in pairs until implantation and then individually in neighboring cages. The temperature (22 ± 1 °C) and humidity (50 ± 20%) in the room were controlled and the room was maintained on a 12-h light/dark cycle (lights on from 07:00 to 19:00 h). In order to motivate animals to collect food rewards, rats were food-restricted up to 85% of their *ad libitum* body weight and subjected to mild water restriction before behavioral sessions (4-12h). The experiments were carried out 5 days a week. During the weekend animals had free access to standard rodent food and unsweetened mineral water.

### Method details

#### Behavioral Setup

The Rat Interactive Foraging Facility (RIFF) arena is described in detail^24^. In short, The RIFF consists of a large circular arena (160 cm in diameter) with 6 equally spaced interaction areas (IAs), each having a water port, a food port (SPECIAL.090-SE v1.0, LaFayette-Campden Instruments, Loughborough, UK), and two free-field loudspeakers (MF1, Tucker Davis Technologies, Alachua, FL, USA), one above each port. Rat behavior is monitored online using video tracking with a monochrome camera (DMK 33G445 GigE, TheImagingSource, Bremen, Germany) mounted above the center of the arena, with a wide-field lens (T3Z3510CS, Computar, North Carolina, US) to capture the whole arena. The ports were controlled and monitored by a commercial software (ABET II, LaFayette-Campden Instruments, Loughborough, UK) and a custom-written MATLAB program (The MathWorks, Inc., Natick, MA, USA). Each IA contained one of 6 flavors of palatable 45 mg food pellets: vanilla, banana, bacon, chocolate, peanut butter and fruit punch (diet AIN-76A: RodTab45MG; RodTabBan45MG; RodTabBcn45MG; RodTabChoc45MG; RodTabPbtr45MG; RodTabFrtP45MG5TUL, TestDiet, Richmond, IN, USA), together with 1 of 4 types of fluids: mineral water, 4% sucrose in mineral water, 0.1% saccharine in mineral water, and 1:1 mixture of 4% sucrose and 0.1 % saccharine solutions (sucrose and saccharin, Sigma-Aldrich, St. Louis, MO, USA).

#### Auditory stimulation

The acoustics of the arena were studied in detail^48^. Absolute sound levels at 0 dB attenuation were about 100dB SPL at 2 kHz, going down to about 85dB SPL at 20 kHz. Sound levels were rather stable as a function of location in the arena (varying by less than 5dB between the center of the arena and the walls at the typical height of the head of the rat) and of loudspeaker (standard deviations across loudspeakers were less than 5dB at the center and less than 8dB at the walls). No correction was applied for this variation.

In active sessions, the rat performed the tasks as described under the Behavioral training section below, and the only sounds presented were the 6 sounds associated with the ports. These sounds were based on the words for “here” or “there” in six different languages (French: la, Italian: qui, English: here, Polish: tam, German: da, and Russian: tut). They were selected from free online recordings (http://shtooka.net), and processed using the STRAIGHT Vocoder^49^ and MATLAB. The spectro-temporal envelope was extracted by STRAIGHT then shifted and stretched along the frequency scale, so that the envelope levels at frequencies 0.1 kHz and 20 kHz of the original sound were shifted to frequencies 1 kHz and 40 kHz with linear interpolation on a logarithmic scale in between. The resulting spectro-temporal envelope of each sound was then superimposed on the original pitch contour. This kept the spectro-temporal modulations of the synthesized sounds as similar as possible to the original ones, while adjusting them to the hearing range of the rats. The first 200 ms of each word was taken as the stimulus, to keep timing consistency between trials. This included the first syllable of each word. To present a sound during the experiment, it was transduced to voltage signals using a sound card (M-16 and MADIface USB, RME Audio Interfaces, Munich, Germany), attenuated by 14 dB (see below), (PA5, Tucker Davis Technologies, Alachua, FL, USA), and then presented through the speakers (MF1, Tucker Davis Technologies, Alachua, FL, USA), driven by stereo power amplifiers (SA1, Tucker Davis Technologies, Alachua, FL, USA).

Passive sessions followed each active session, and included recordings of additional stimuli used to characterize the auditory responsiveness at the electrode location. Here we present the responses only to the task-relevant stimuli. The word-like sounds were presented in a pseudorandom order and at a rate of 1 Hz at 14dB attenuation from maximal sound level (same as for the behavioral session). Each sound was repeated 20 times, for a total of 120 presentations of the word-like stimuli during the passive session. The sounds were presented in a small enclosure inside the arena.

### Behavioral task

#### Training

Rats (N=4) were habituated to the RIFF in two 12-hour night sessions. During habituation, the RIFF was logically divided into 6 equal sectors centered on the IAs (“pie slices”), and each of these sectors was associated with a specific word-like stimulus that could be activated according to the behavior of the rat (a different stimulus for each IA, sound mapping was the same in 3 rats and shifted counter-clockwise by 2 IAs for one of the rats). Rats were placed in the RIFF and were able to explore the arena freely. Whenever the rat entered an active sector the associated sound was presented from the two speakers of the IA in that sector. The sound repeated every 2 s until the rat poked in either of the ports of the IA and got a reward, with a timeout after 2 minutes. Before the first poke all sectors were activated when the rat entered them. After a nose poke or a timeout, the current sector and its two neighbors became inactive but the three other sectors became active, so that the rat had to cross to one of the three sectors on the opposite side of the RIFF in order to initiate a new interaction. Following habituation, the rats were exposed to increasingly stricter versions of the main task (see below, and Fig. 1a) in 12 hour overnight sessions. For the first 6 training sessions the rats were allowed to poke in more than one port before trial termination (10 pokes in the first and second sessions, decreased to 6 and 4 for one session each, 2 pokes for 2 sessions, then down to 1 for the rest of the experiment). In addition, the requirements for initiating a trial became stricter: on the first training session, the center had a radius of 50 cm, on the 2nd session of 40 cm, and from the 3rd session on the center had its final radius of 30 cm. Then for 2 sessions the next target was always one of the three sectors on the opposite side of the RIFF, to encourage the rats to cross the center. The rats were exposed to the full main task after this training procedure.

#### Main task

The rats initiated trials by moving into a central circular area (radius of 30 cm, Fig. 1a). One of the 6 IAs was randomly selected by the program and between 500-700 ms after the detection of a crossing into the central area, its associated sound was presented from its two speakers. The same sound was presented every 2 s (up to 10 times, maximal trial length 20 s), or until the rat poked in any port. A poke in one of the two ports of the IA from which the sound was played resulted in a reward (food or water, according to the poked port). A poke in a wrong port, or 20 s without a poke, resulted in the termination of the trial (see Fig. 1a). There was no penalty for unsuccessful trials, so that rats could immediately initiate a new trial by moving once more into the central area.

#### Control 1: pure localization

For the localization control, one of the 6 sounds was randomly selected and for a batch of 40 trials, that sound was played from the two speakers of any of the selected target locations. In consequence, the rats could only use the location of the speakers as a cue for reward. For each batch of 40 trials, a different sound was selected. These batches were interleaved with batches of main task trials (40 trials per batch).

#### Control 2: pure discrimination

In this task each target sound was played from all speakers simultaneously, and its identity determined the selected IA (6-way arbitrary association task). The task turned out to be very hard for the rats. In order to make it easier an adaptive staircase procedure was used. The target speaker played the sound always at 20 dB attenuation while the non-target speakers played the sound at a different sound level (the same for all speakers), varying from 50 dB attenuation up to 20 dB attenuation (in 3 dB jumps). The attenuation for the non-target speakers was set, trial by trial, based on the performance of the rat in the last 8 trials. If 6 or more trials among the last 8 were successful, the sound level of the non-target speakers was increased by 3 dB (attenuation decreased by 3 dB, making their sound level more similar to the target speaker); if 6 or more trials among the last 8 were unsuccessful, the sound level of the non-target speakers was decreased by 3dB; otherwise the sound level of the non-target speakers was kept unchanged. These blocks lasted whole sessions. The staircase procedure ensured a success rate averaging about 50%.

#### Procedure

Rats were trained for 2 nights on the habituation paradigm, then at least 5 weeks on the main task before implantation of the base of the electrodes (see Jankowski et al. 2023 for details of the surgical procedures). Three to five weeks after base implantation the rats were implanted with the silicon probes and were switched from 12 hour overnight sessions to 3 hour long day sessions. After electrode implantation, neuronal responses were recorded for 10-15 daily sessions of the main task and 10-15 sessions for each control (see Behavioral task section). Then main task and control sessions were presented for 3-5 consecutive days in a cycle until the end of the experiment. The rats performed the main task in the arena for 2.5 hours (‘active session’), following which the rats were placed in a smaller enclosure for a passive listening session that lasted about 30 minutes (see above, Auditory Stimulation).

#### Electrophysiology and recordings

The electrophysiological methods are described in detail^24^. In short, a 64 channel neural logger (RatLog-64, Deuteron Technologies, Jerusalem, Israel) was used throughout the experiment. Rats were chronically implanted with 32-channel silicon probes (ASSY-116_E-2, Cambridge Neurotech, UK) in the auditory cortex. The latencies of the unit responses to broadband noise bursts were the expected in the core auditory cortex (A1 or AAF; 13ms (10.5-15.5ms), median (IQR), N=819; all units with significant responses to broadband noise). Neural signals were recorded in reference to a ground placed in the frontal bone. For the logger, the analog bandpass filter was set to 10-7000 Hz. Recording sessions were performed five days a week. Spiking activity was screened immediately following each recording session. The electrodes were kept in the same position as long as spiking activity was detected on many contacts. When signals deteriorated, the animals were briefly sedated with sevoflurane and the electrodes were lowered in steps of 25, 50 or 100 µm into the brain tissue. Electrodes were moved down every 1-7 days. The number of recording sessions were 47 (20 main task, 13 localization control, 14 discrimination control; rat 4), 63 (26 main task, 19 localization control, 18 discrimination control; rat 5), 51 (19 main task, 18 localization control, 14 discrimination control; rat 6) and 48 (15 main task, 16 localization control, 17 discrimination control; rat 8). Recordings were terminated when neural responses were lost on all electrodes.

#### Data processing

Initial data processing was performed using the analysis pipeline and the post processing as described^24^, and included spike sorting and feature extraction from the video stream. The analysis of the responses to the task-relevant sounds was performed on data discretized into 1 ms bins. For trial-level analysis, the data was discretized into 100 ms bins.

### Quantification and statistical analysis

Statistical analysis was performed using MATLAB (The MathWorks, Inc., Natick, MA, USA). To assess responsiveness to auditory stimuli of all units in passive and active conditions we used an inhomogeneous Poisson likelihood ratio test^50^. We assumed that the on-going firing rate (taken during 200 ms preceding the stimulus) was a homogeneous Poisson process, and tested it against rates that were computed in 20 windows of 10 ms (a total of 200 ms) following stimulus onset. The logarithms of the probability ratios were summed to form the log-likelihood ratio (LL). We tested 2LL against a χ^2^(20) distribution. Responses were considered significant when p<0.05. To compare the sound driven responses (0-200 ms after sound onset, throughout the duration of the sound) in the passive and active conditions we used linear mixed effects models for the mean firing rate in the appropriate time window of each unit as a function of condition. The Wilcoxon notation for the model was fr∼condition+(condition|unit:rat)+(sound-1|unit:rat)’. Here fr was the mean firing rate, condition was active or passive, and all other variables had their natural interpretations. The random factors were used to account for the correlated variability within each unit. To compare on-going activity (−200-0 ms before sound onset) in the passive and active conditions we used linear mixed effects models for the mean firing rate in the appropriate time window of each unit as a function of condition (Wilcoxon notation: fr∼condition+(condition|unit:rat)’).

The variability index measured the discrepancy between the responses averaged over all sounds and the responses to each of the sounds individually. It was computed using the following formula:

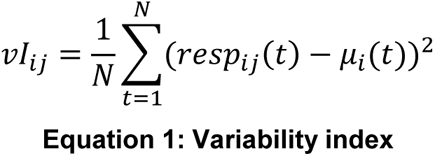

Where i is the rat index (rat 4, 5, 6 or 8), j is the sound index (one of 6 sounds), resp_ij_(t) was the response (averaged over all units) to sound j in rat i at time t, μ_i_(t) was the response averaged over all units and all sounds in rat i at time t. The squared differences were average over the stimulus presentation time, from sound onset to sound offset, a total of N=200 ms. There were a few hundred sound presentations in each active session (distributed over the 6 task-relevant sounds), but only 20 presentations of each task-relevant sound in the passive condition. While the difference in the number of averaged trials could affect the variability of this measure, we didn’t compensate for that. Indeed, we expected such additional variability to inflate the variability index in the passive condition, but the variability index tended to be larger in the active condition. Thus, the comparison is, if anything, conservative.

To calculate the mutual information (MI) between the sound identity and the responses in both active and passive conditions we used principal component analysis on the single trial responses of each unit (using the MATLAB function ‘svd’), and then calculated the MI between the projections on the first singular component of the single-trial responses and the sound identity (one of 6 sounds), using a procedure^51^ that compensated for the finite sample bias.

To study the relevance of acoustic and non-acoustic factors in determining the firing rates, linear mixed effects models (MATLAB function ‘fitlme’) were fitted to the firing rates (defined as spike counts per 100ms bins) of the individual units as a function of each one of several auditory or behaviorally relevant parameters (each one is referred to as the main explanatory variable of their respective model). These models were fitted to the data recorded from each rat separately, but included all units from all recording sessions of each rat. A detailed description of one of these models is provided here. The models differed only in their main explanatory variable. All models are described in Table 1.

**Table 1:**
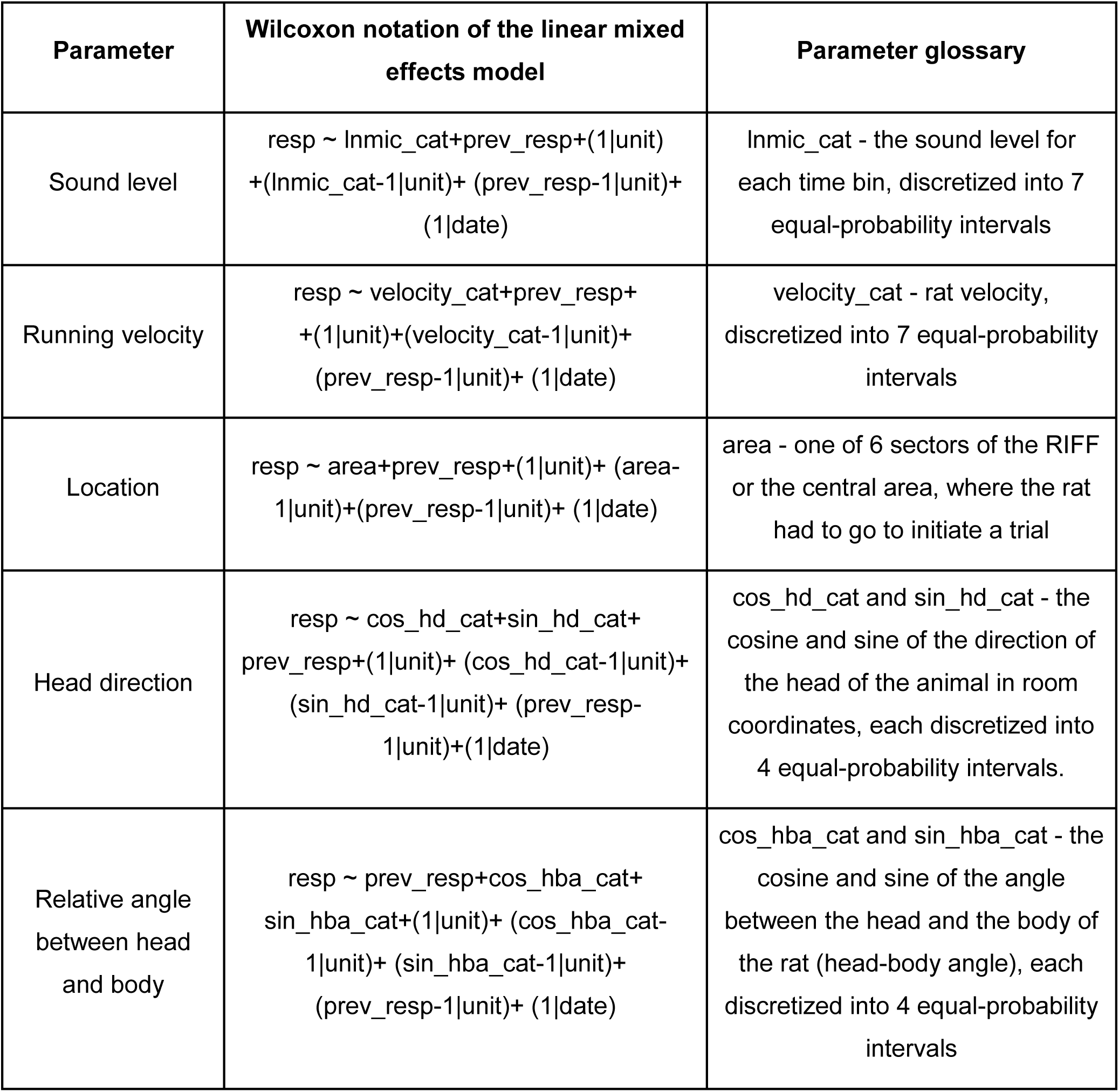
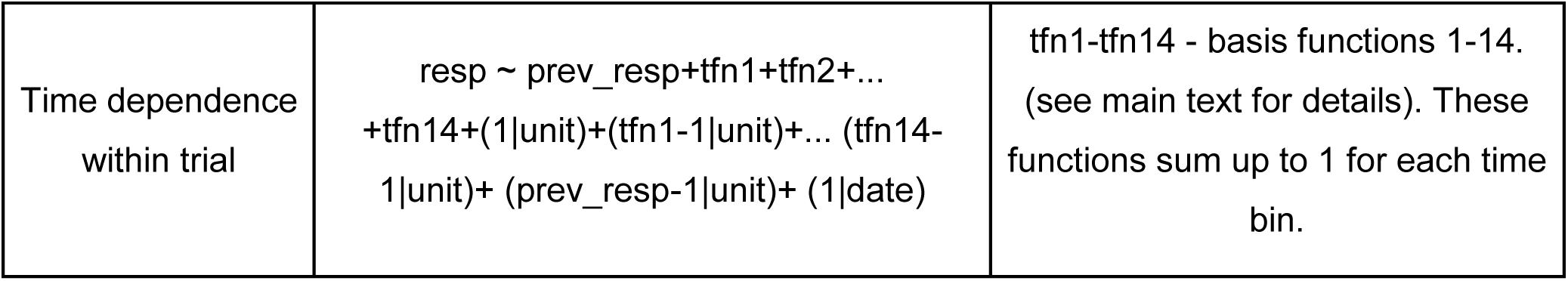
Wilcoxon notation of each LME model used for data analysis.

The model described here used the sound level at the microphone on the headstage as the main explanatory variable. It modeled the firing rate of the unit as depending on previous firing rates as well as on the sound level (see Table 1 for its Wilkinson notation):

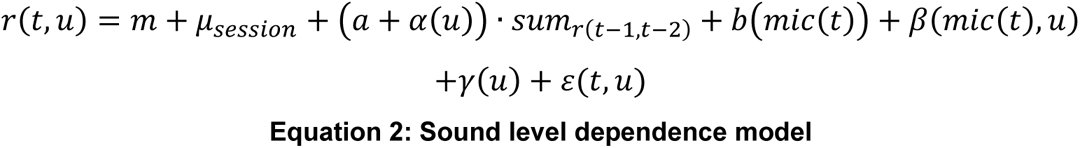

Here r(t,u) is the spike count of unit u at time t. The explanatory variables always included the sum of the spike counts in the previous two bins, *sum*_*r*(*t*−1,*t*−2)_ = *r*(*t* − 1, *u*) + *r*(*t* − 2, *u*), since firing rates tended to show slow changes and so were correlated in nearby times. The sound level was coded into one of 7 equal-probability intervals (whose index was used as a categorical variable in the model), so that the effect of sound level on the firing rate was modeled as a nonlinear function that was constant on each of these intervals. The coefficients of the two explanatory variables, previous spike count and sound level were separated into ‘main effects’ (denoted by *m*, *a*, and *b*(*i*) in Eq. 2) that accounted for the moment-by-moment firing rate averaged over all units, and ‘random effects’ which accounted for the deviations of the unit-specific coefficients from the respective main effect. One additional random effect was introduced to account for differences in mean firing rates between recording sessions (denoted by *μ*_*session*_ in Eq. 2). The unit-specific random effects are denoted by α (coefficient of the firing rate in the preceding two time bins), β (for the sound-level dependence) and γ (unit specific intercept). These deviations were regularized by estimating them under the assumption that they were Gaussian with mean 0 and a covariance matrix that was estimated from the data by maximizing the likelihood^52^. The unit-specific coefficients were estimated as the Best Linear Unbiased Predictors (BLUPs^53^) given the fixed factors and the covariance matrices of the random factors.

All models had the same basic structure, except for the main explanatory variable. We estimated models for the dependence of the firing rates of the units on the following variables: sound level (as described above in detail); running velocity (coded as the index of one of 7 equal-probability intervals); location (coded as one of 7 areas: one of 6 equal sectors centered around an IA, or the circular center area); head direction (coded as 4 equal probability categories for the sine and 4 equal probability categories for the cosine of the angle in room coordinates); the angle between the head and the body of the rat (coded in the same way as head direction); Lastly, models with explicit time-dependence were estimated. The time within trial was coded using 14 triangular basis functions of width 700 ms (7 time bins), with an overlap of half width. These functions covered a total of 4 seconds, starting from 2 s before center crossing and ending at 2 s after center crossing, and spanned the space of all continuous functions that are linear over each of the 200 ms intervals covering this time range. The time range included as much time of the trial as possible while keeping the number of degrees of freedom of the model comparable to that of the other models. The Wilcoxon notation for each model is shown in Table 1, where ‘resp’ denotes the spike count in 100ms bin, ‘prev_resp’ denotes the sum of the firing rates in the two time bins preceding each bin, ‘unit’ denotes the identity of each of the units recorded in the rat, and ‘date’ denotes the individual sessions.

The models were compared to each other using the Akaike Information Criterion (AIC). This criterion is an estimate of the generalization error of the model, thus allowing for direct comparison between several models of the same data even when they are not nested within each other^54^. The AIC takes into account the number of parameters (k) in each model:

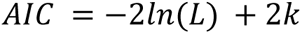

Where L is the likelihood of the model. Thus, a smaller AIC indicates a better fit to the data. The linear mixed effects models also provide fits for each unit individually. The log likelihood, L_unit_, for each unit separately in each model was calculated as follows:

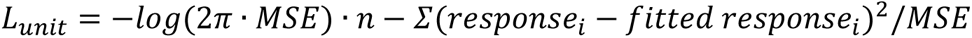

Where n is the total number of entries in the data table containing data for the specific unit, the sum is over all of these entries in the table that are within the relevant time range, *response*_*i*_ is the firing rate, *fitted response*_*i*_ is the fit of the model, and MSE is the mean squared error of the model.

### Model

For modeling the cortical responses, the peripheral representations of the task-relevant sounds were computed using the Bruce auditory-nerve model^33^ implemented in the auditory modeling toolbox^55^ (AMT) version 1.2. We simulated 32 frequency channels, equally spaced on a logarithmic axis between 1 kHz and 40 kHz. These were further processed to mimic the phasic responses of forebrain neurons, by first low-pass filtering the time course of each frequency channel (with a cutoff frequency of 70 Hz), followed by differentiation and half-wave rectification. The cortical model^30^ was modified with respect to its original form in two ways only. First, the original model was designed for pure tone input only. The simulation was therefore adjusted to use the computed simultaneous input on all frequency channels. The model had 32 frequency columns, and each model neuron was driven by the frequency channel corresponding to its best frequency.

The second modification consisted of modifying the baseline firing rate of the neurons, in order to simulate responses in the active and in the passive conditions. In the original model^30^, the baseline inputs to the neurons in each column ranged between −10 and 10 sp/s. To simulate changes in these baseline rates, these ranges were modified: the lowest rates were between −6 and 6 sp/s, and the highest were between −14 and 14 sp/s, at a resolution of 1 sp/s (9 values). For simulating active and passive conditions, each of the conditions was compared with that of another condition where the range of the baseline firing rates was higher by 3 sp/s, corresponding to a change of about 30% in the baseline firing rates.

## Author contributions

IN conceived and supervised the project. AP designed and wrote the software for implementing the behavioral paradigm with a contribution of JN and AK. MMJ designed and developed the chronic neural implants. AP and MMJ collected data. JN, AK, and AP developed the basic analysis pipeline and spike sorting of the data. AP analyzed the data presented in this paper. AP and IN wrote and approved the final manuscript.

## Acknowledgements

We thank Mousa Karayanni for helping with the data collection.

This work was supported by AdERC grant GA-340063 (project RATLAND), by F.I.R.S.T. grant no. 1075/2013 from the Israel Science Foundation. JN was supported by a DFG Research Fellowship (ref. NI 2012/1-1; project number 442068558). IN holds the Milton and Brindell Gottlieb Chair in Brain Sciences.

## Declaration of interests

The authors declare no competing interests

## Notes

https://doi.org/10.6084/m9.figshare.26310223

### Competing Interest Statement

The authors have declared no competing interest.

