## Supplementary Information for "Task-related activity in auditory cortex enhances sound representation"

### Supplementary Figures S1-S5
### Supplementary Tables S1-S10

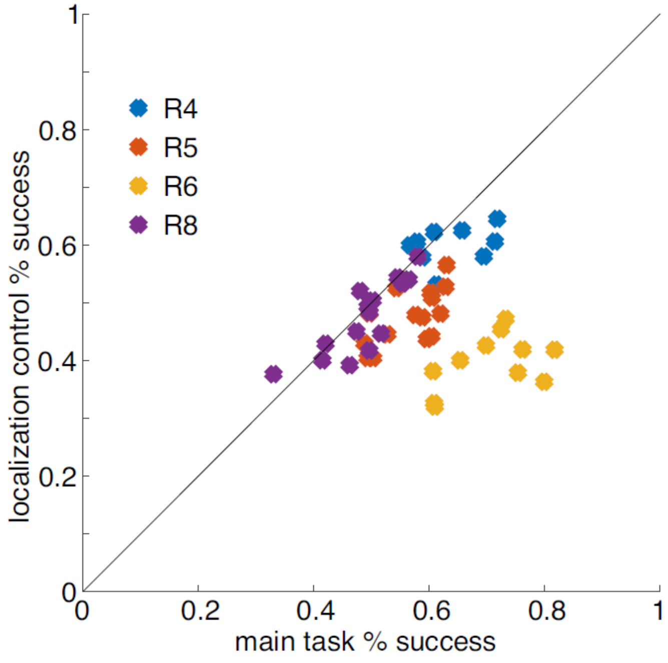


**Figure S1. Success rate in the main task blocks compared to the localization control blocks.** Each point corresponds to one session. Each color corresponds to a different rat.

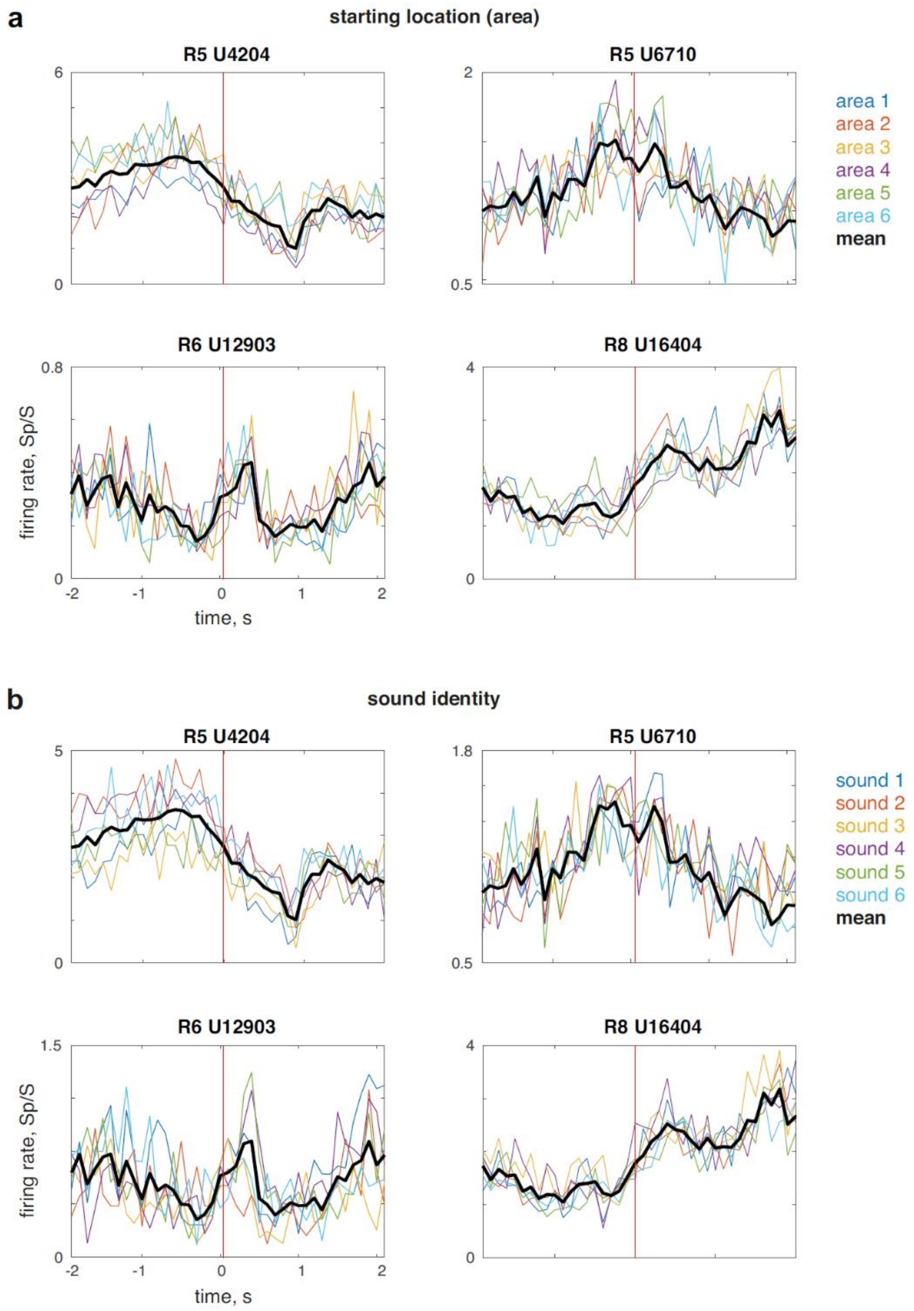


**Figure S2. Firing rates were independent of location at trial initiation or sound presentation after center crossing.** a. Mean firing rate around center-crossing (time 0) divided by the port in which it got the previous reward. This port is the point from which the rat initiated the next trial. Thick black line is the overall mean response throughout all trials. Red vertical line indicates trial crossing. b. Same as (a) but for the 6 different sounds presented to the rat.

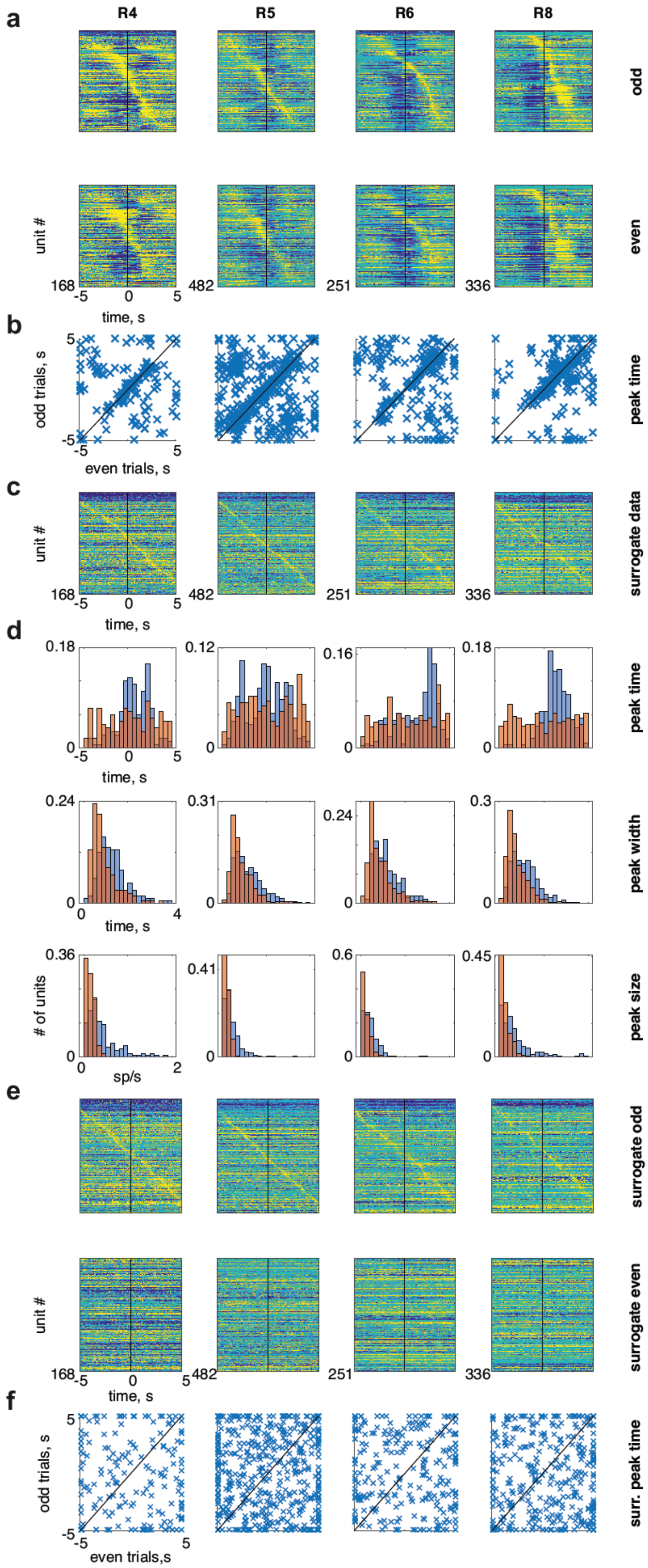
a
R4
R5
R6
R8
odd
even
unit #
168
482
251
336
-5
0
5
time, s
b
odd trials, s
even trials, s
peak time
c
surrogate data
d
0.18
0.12
0.16
peak time
0.24
0.31
0.3
peak width
0.36
0.41
0.6
0.45
### of units
sp/s
peak size
4
2
e
surrogate odd
surrogate even
f
even trials,s
surr. peak time

**Figure S3. Large slow firing rate modulations and controls in all rats.** Data from one rat in each column. a. Top: normalized mean firing rate of all units averaged over odd trials, sorted by the time of the maximum firing rates. Bottom: normalized mean firing rate of all units averaged over the even trials, same order as top row. These are the same plots as in Fig. 4 b. The time of the maximal firing rate in the averages over even vs. odd trials for each unit. c. Mean firing rates around uniformly distributed surrogate center-crossing times d. Distribution of peak time (top), width (middle) and size (bottom) of the maximal firing rates of the real (blue) and surrogate (red) data. e. Top: normalized mean firing rate of all units averaged over odd trials in the surrogate data, sorted by time of the maximum firing rates. Bottom: normalized mean firing rate of all units averaged over the even trials, same order as top row. f. The time of the maximal firing rate in the averages over even vs. odd trials for each unit in the surrogate data.

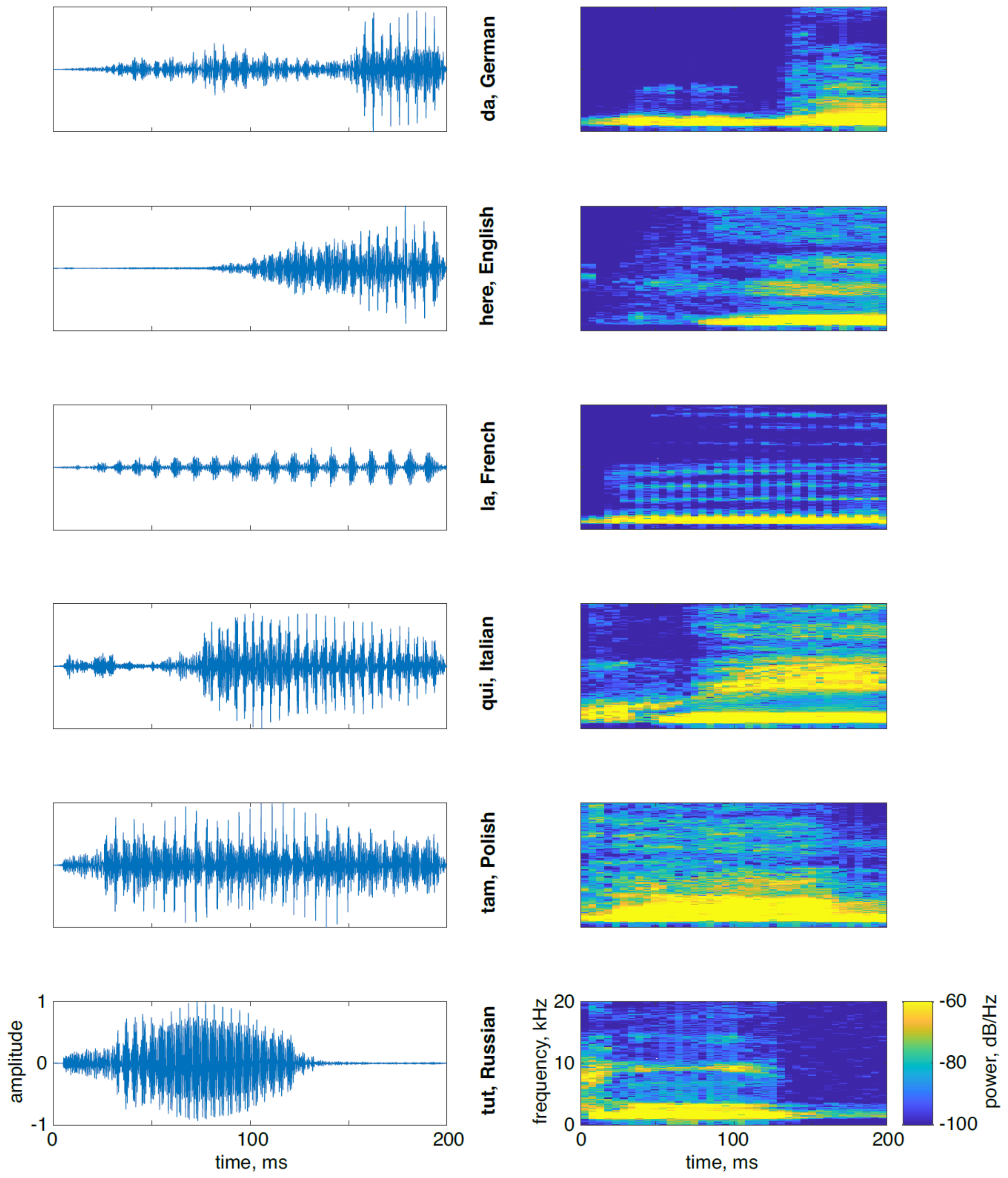


**Figure S4. Oscillograms (left) and power-spectra (right) of the task-relevant sounds.**

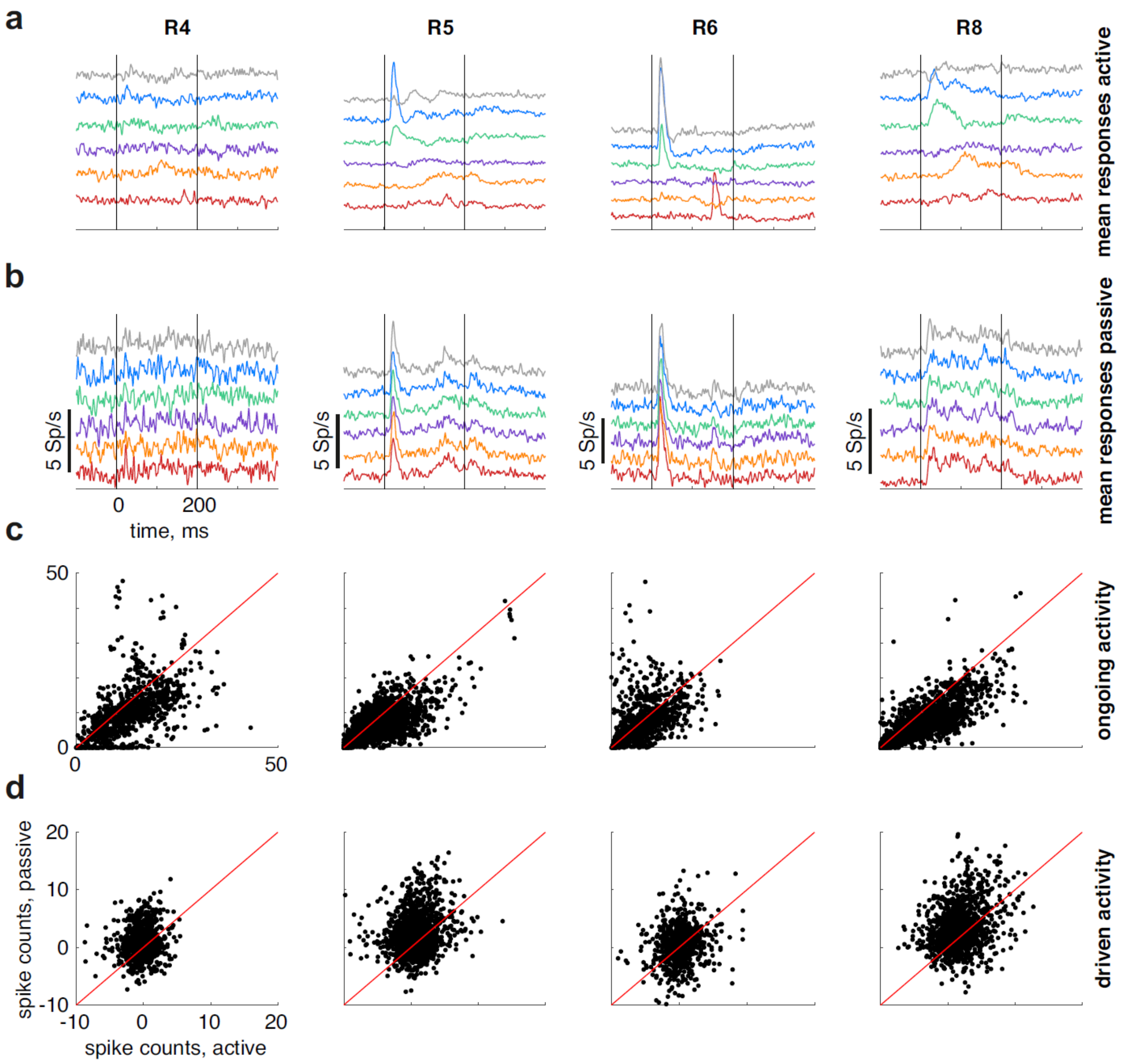


**Figure S5. Responses to the task-relevant sounds in all rats.** Data from one rat in each column. a. Mean responses averaged over all units to the task-relevant sounds in the active sessions. Each sound is displayed in a different color. b. Same for the passive sessions. c. Mean spike counts of the units of the on-going activity in the 200 ms preceding sound onset during active compared to passive sessions. Each dot is one unit. d. Mean spike counts of the units during sound presentation (0-200 ms after sound onset) during active compared to passive sessions. Each dot is one unit.

**Table S1:** comparison between locations of peaks in odd and even trials

| | $\chi^2(16)$ | p-value |
|---|---|---|
| R4 | 98.2 | $7.6*10^{-13}$ |
| R5 | 250 | $3.6*10^{-44}$ |
| R6 | 158 | $1.4*10^{-25}$ |
| R8 | 288 | $5.9*10^{-52}$ |

**Table S2:** comparison of location, height and width of the peaks in the real and surrogate data

| | **Location of maxima** | | **Width of maxima** | | **Size of maxima** | |
|---|---|---|---|---|---|---|
| | D | p-value | t | p-value | t | p-value |
| R4 | D(168,168) = 0.23 | $2.9*10^{-4}$ | t(334) = 6.04 | $4.0*10^{-9}$ | t(334) = 9.27 | $2.4*10^{-18}$ |
| R5 | D(482,482) = 0.17 | $1.9*10^{-6}$ | t(962) = 8.59 | $3.2*10^{-17}$ | t(962) = 11.4 | $2.3*10^{-28}$ |
| R6 | D(251,251) = 0.22 | $1.4*10^{-5}$ | t(500) = 5.91 | $6.3*10^{-9}$ | t(500) = 7.67 | $9.1*10^{-14}$ |
| R8 | D(336,336) = 0.39 | $1.1*10^{-22}$ | t(670) = 8.79 | $1.2*10^{-17}$ | t(670) = 9.72 | $5.6*10^{-21}$ |

**Table S3:** comparison between location of peaks in odd and even arbitrarily chosen 10 second periods

| | $\chi^2(16)$ | p-value |
|---|---|---|
| R4 | 11.9 | 0.75 |
| R5 | 8.26 | 0.94 |
| R6 | 8.40 | 0.94 |
| R8 | 20.2 | 0.21 |

**Table S4:** AIC measurements in all rats

AIC - min(AIC) = $10^4$ *

| | **Head body angle** | **Head direction** | **Location** | **Velocity** | **Time dependence** | **Sound level** |
|---|---|---|---|---|---|---|
| R4 | 1.96 | 1.89 | 1.79 | 1.88 | 1.26 | 0 |
| R5 | 4.75 | 4.29 | 3.42 | 3.35 | 2.38 | 0 |
| R6 | 2.98 | 2.67 | 1.69 | 0.73 | 1.02 | 0 |
| R8 | 3.79 | 3.62 | 2.87 | 2.14 | 0 | 0.99 |

**Table S5:** On-going firing rates in active compared to passive conditions, linear mixed effects model, Wilcoxon notation: fr~condition+(condition|sound:unit:rat)+(sound-1|unit:rat)

| Term | F(1,12262) | p-value |
|---|---|---|
| intercept | 433 | $1.3*10^{-94}$ |
| condition | 238 | $3.5*10^{-53}$ |

**Table S6:** Firing rates in the 200ms following sound onset  in active compared to passive conditions, linear mixed effects model, Wilcoxon notation: ‘fr~condition+(condition|unit:rat)

| Term | F(1,12262) | p-value |
|---|---|---|
| intercept | 1521 | $9.9*10^{-314}$ |
| condition | 241 | $6.2*10^{-54}$ |

**Table S7:** Mutual information between sound identity and the responses, linear mixed effects model, Wilcoxon notation ‘MI~condition+(1|rat)+(1|unit:rat)

| Term | F(1,1700) | p-value |
|---|---|---|
| intercept | 633 | $5.2*10^{-119}$ |
| condition | 107 | $2.2*10^{-24}$ |

**Table S8:** Comparison between slow firing rate events and sound-driven responses, linear mixed effects model, Wilcoxon notation: 'resp~type+(type|rat)', type was sound-driven response or slow firing rate event

| Term | F(1,2472) | p-value |
|---|---|---|
| intercept | 0.22 | 0.64 |
| condition | 41.8 | $1.2*10^{-10}$ |

**Table S9:** Comparison of the mean firing rate of a unit as a function of one of three conditions: ongoing activity just before sound onset in the active condition; the ongoing activity just before sound onset in the passive condition; and the minimal firing rates within the +-5 s window around center-crossing, with condition within rat as a random factor; Linear mixed effects model, Wilcoxon notation: 'resp~condition+(condition|rat)'

| Term | F(1,3114) | p-value |
|---|---|---|
| intercept | 41.6 | $1.3*10^{-10}$ |
| condition | 16.5 | $7.3*10^{-8}$ |

**Table S10:** Correlation between minimal active and ongoing passive.

| | $R^2$ | p-value |
|---|---|---|
| R4 | 0.58 | $3.0*10^{-16}$ |
| R5 | 0.82 | $2.4*10^{-92}$ |
| R6 | 0.52 | $1.2*10^{-12}$ |
| R8 | 0.75 | $7.8*10^{-61}$ |
